# Warmth shifts rather than speeds transcriptomic ageing in the malaria mosquito *Anopheles gambiae*

**DOI:** 10.64898/2026.10.13.758761

**Authors:** Krishnendu Sinha, Nabanita Ghosh, Sharmistha Banerjee

## Abstract

Only older mosquitoes transmit malaria, so how warming changes mosquito ageing matters for disease. Warmer temperatures are known to make mosquitoes age faster, but “faster” can mean two different things: the ageing clock ticks more quickly, or it is simply set forward. We distinguished these by reanalysing 115 whole-body RNA-sequencing libraries from female *Anopheles gambiae* kept at 27, 30 or 32 °C and sampled 1–15 days after emergence. A transcriptomic clock trained on 27 °C mosquitoes (error *±*0.9 days) read 32 °C mosquitoes as 2.4 days older (95% CI 1.9–2.9) and 30 °C mosquitoes as 1.1 days older. The gap opened by day 5 and then stopped growing: between days 5 and 15, warmth added 0.01 extra days per day (95% CI *−*0.16 to 0.19), well below the 0.29–0.45 expected if the clock ran faster. Genome-wide, warmth advanced genes that change late in life, notably melanisation and digestive genes, but moved early-maturation genes the opposite way. Many responses bent at 30 °C, and immune genes held until 30 °C before falling. Warmth therefore shifts mosquitoes to an older transcriptional state without accelerating its pace.

## Introduction

A mosquito’s body is as warm as the air around it, so a few degrees of environmental warming changes the speed of almost every process inside it. For malaria this matters in a particular way. After a female *Anopheles gambiae* takes an infected blood meal, the parasite needs roughly ten days or more to develop before it reaches the salivary glands, and the mosquito can only transmit if it survives that long. Transmission therefore depends heavily on how many mosquitoes live into old age, and on how well their bodies, including their immune systems, function when they get there. Because the parasite’s development and the mosquito’s survival both respond to temperature, and not in a straight line, transmission peaks at intermediate temperatures and declines at the warm end [1].

Several lines of evidence suggest that warmth makes mosquitoes age faster. *A. gambiae* reared and kept at higher temperatures survive for shorter periods [2]. Their melanisation immune response, which encases pathogens in melanin, weakens with age, and it begins to weaken earlier at 30 °C and 32 °C than at 27 °C [3]. Using the RNA-sequencing data reanalysed here, Martin and colleagues showed that warmer temperature modifies the age-dependent changes in gene expression that involve immunity, metabolism and DNA repair, and concluded that warmth accelerates senescence [4].

“Accelerated ageing” can, however, mean two different things, and the difference has practical consequences. Warmth could make the ageing clock *tick faster*, so that each day at 32 °C counts for more than a day at 27 °C and warm mosquitoes drift ever further ahead as they age. Alternatively, warmth could *set the clock forward*: warm mosquitoes would reach an older-looking state early in adult life, and from then on age at the normal pace (Fig. 1b). The two ideas predict different futures. A faster clock compounds over a mosquito’s lifetime, whereas a forward shift does not. A comparable distinction transformed the study of dietary restriction in *Drosophila*, which was shown to lower the immediate risk of death rather than slow the accumulation of ageing damage [5]. Telling the two apart requires measurements at several ages, which most thermal studies lack but the present data set provides.

**Figure 1:**
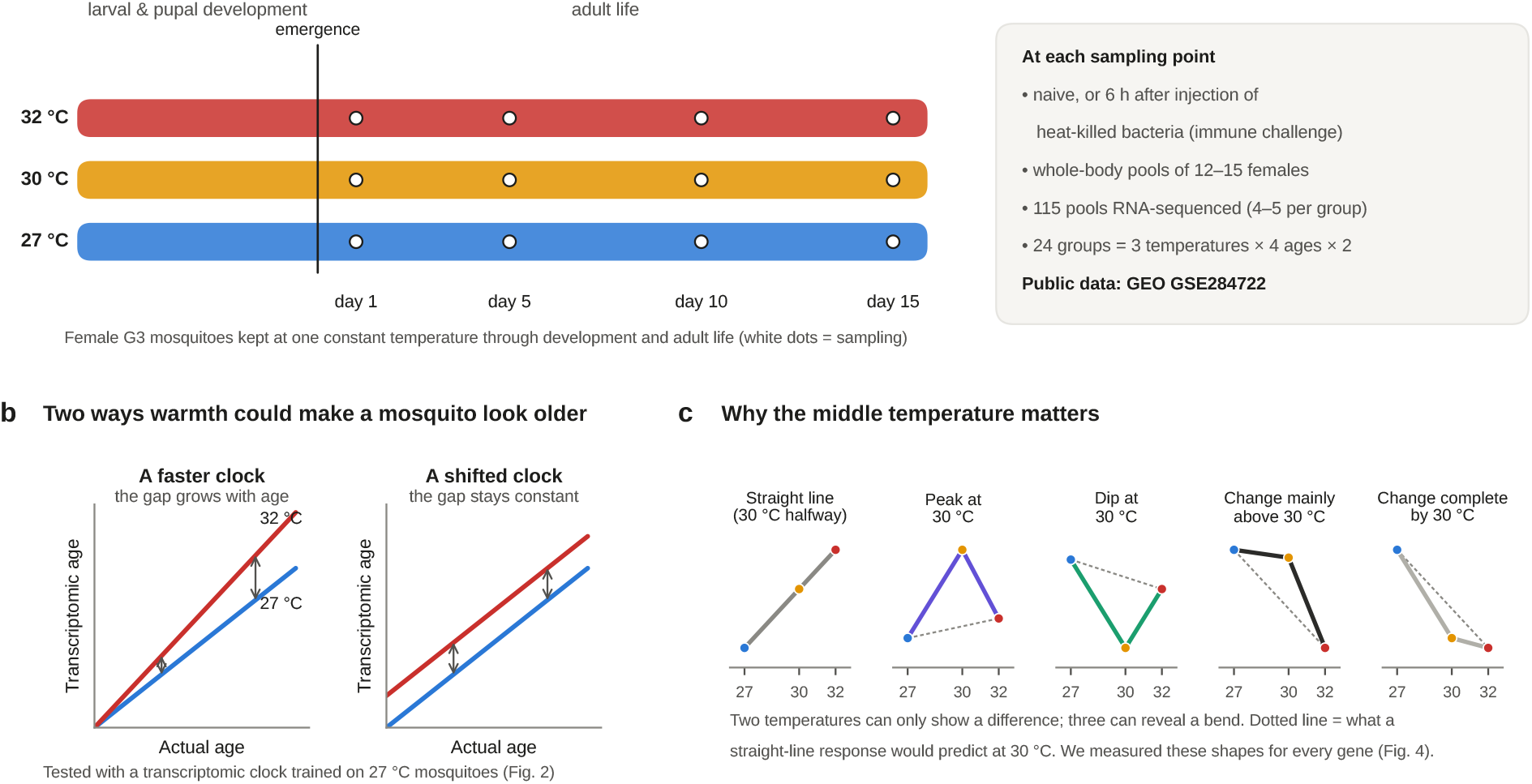
Design and the two questions it can answer. **a**, Female *A. gambiae* (G3 strain) were held at 27, 30 or 32 °C through development and adult life and sampled 1, 5, 10 and 15 days after emergence, either untreated (naive) or 6 h after injection of heat-killed *Escherichia coli* (immune challenge). Each RNA-sequencing library is a pool of 12–15 whole females; 115 pools passed the source study’s quality control. **b**, Two ways warmth could make a mosquito look older. If the clock ticks faster, the gap between warm and cool mosquitoes grows with age; if it is shifted forward, the gap stays constant. **c**, A third temperature reveals whether a response bends. The dotted line shows where 30 °C would sit if the response were a straight line between 27 and 32 °C.

A second gap concerns the shape of thermal responses. Most experiments compare two temperatures, which can reveal a difference but not whether a response rises steadily, peaks at an intermediate temperature or changes abruptly beyond a threshold. Physiological traits of ectotherms, including mosquito immune function, often respond to temperature in curved, non-monotonic ways [1, 6]. With three temperatures, the middle one shows whether a response is a straight line or bends (Fig. 1c).

Here we reanalysed the public 27–30–32 °C time series of Martin *et al*. [4] to ask four questions. First, does warmth make mosquitoes look older because their transcriptomic clock runs faster, or because it is shifted forward? Second, which parts of the ageing programme are advanced, and do any move the other way? Third, how often does the 30 °C state differ from a halfway point between 27 and 32 °C? Fourth, do RNA processing, repeat-derived RNA and between-sample consistency change in step with gene expression? We find that warmth shifts transcriptomic age forward by about 2.4 days at 32 °C without changing its pace, that the shift reflects late-life rather than early-maturation biology, and that many responses, including those of immune genes, change mainly between 30 and 32 °C.

## Results

### A transcriptomic clock reads mosquito age to within a day

The data comprised 115 pools spread over 24 groups (3 temperatures *×* 4 ages *×* 2 immune states, 4–5 pools each; Fig. 1a), and 11,151 genes were expressed well enough to analyse. To measure transcriptomic age we built a simple clock: a regression that predicts a pool’s age in days from the expression of the 200 genes most strongly correlated with age in naive 27 °C mosquitoes, compressed into three principal components (Methods). The clock was always tested on mosquitoes it had not seen. The experiment was run in six cohorts, and for each cohort the clock was rebuilt from the other five, so that gene selection and model fitting never used the cohort being predicted.

Tested this way, the clock read the age of naive 27 °C mosquitoes with a mean absolute error of 0.90 days (root-mean-square error 1.20 days; *R*^2^ = 0.95; *n* = 18; Fig. 2a). It is therefore a good ruler for transcriptomic age within this experiment. It is not a universal mosquito age clock, and it says nothing directly about remaining lifespan.

**Figure 2:**
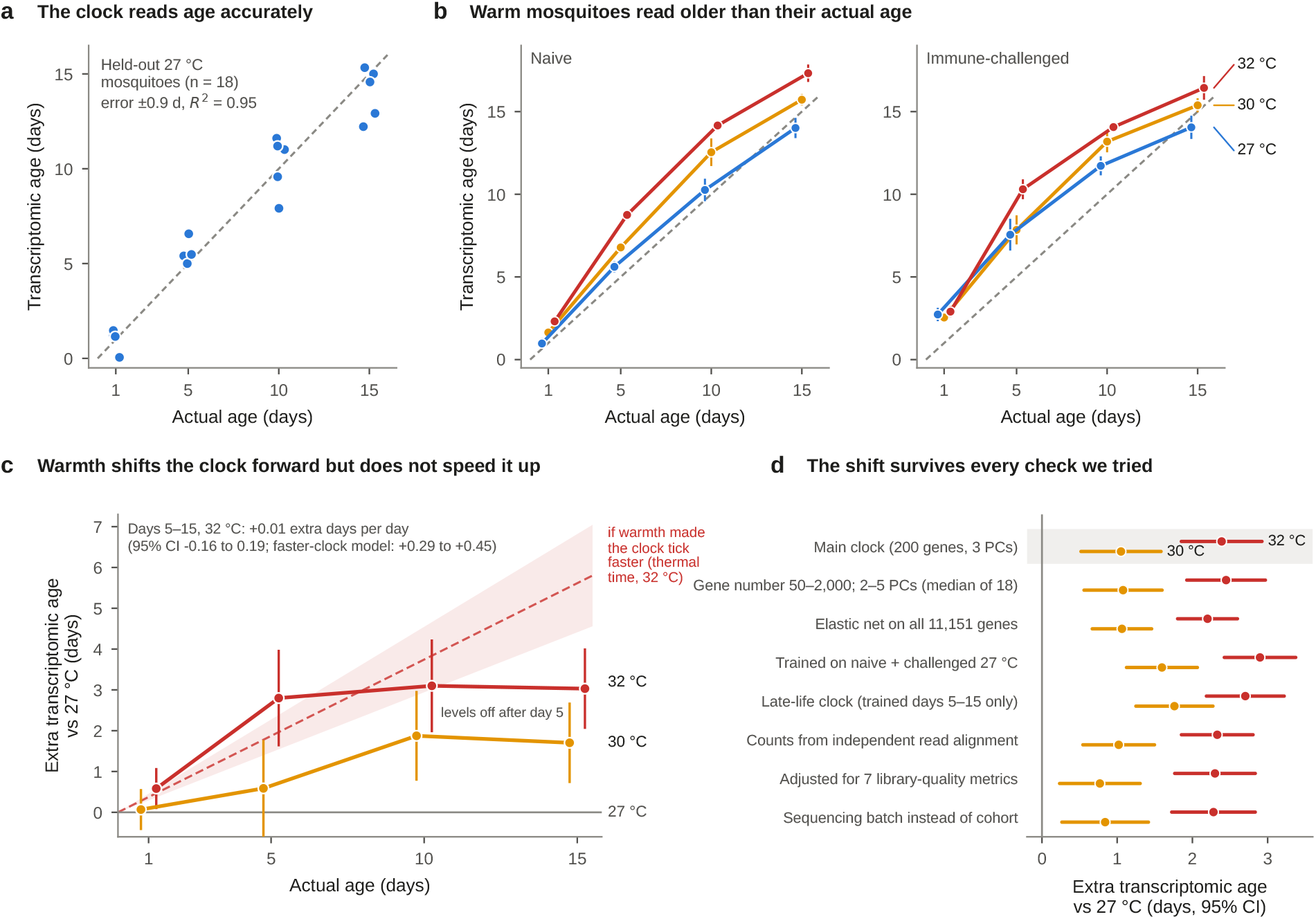
Warmth shifts the transcriptomic clock forward but does not make it run faster. **a**, Clock accuracy in naive 27 °C mosquitoes, each predicted by a clock that never saw its cohort. Dashed line, perfect prediction. **b**, Transcriptomic age against actual age for each temperature (means *±* s.e. across pools). **c**, Extra transcriptomic age relative to 27 °C mosquitoes of the same age (pooled over immune state, cohort-adjusted, 95% CI). The shaded band shows what a faster clock would produce if transcriptomic age accumulated in proportion to thermal time above a lower threshold of 10–16 °C; the dashed line is its midpoint. **d**, The 32 °C (red) and 30 °C (amber) shifts under alternative clock constructions and data-processing choices (95% CI). The grey row is the clock used throughout. All 29 clock constructions are shown in Supplementary Fig. S1b.

### Warm mosquitoes read older than their actual age

Applied to all 115 pools, the clock read mosquitoes kept at 30 °C and 32 °C as older than 27 °C mosquitoes of the same age (Fig. 2b). Averaged across ages and immune states, and accounting for cohort, the extra transcriptomic age was 2.39 days at 32 °C (95% CI 1.85–2.92; *P* = 3 *×* 10^*-14*^) and 1.05 days at 30 °C (0.52–1.58; *P* = 2 *×* 10^*-4*^). Terms involving temperature as a whole improved the model substantially (*F*_16_,_86_ = 6.89, *P* = 8 *×* 10^*-10*^), whereas there was no evidence that the temperature effect depended jointly on age and immune state (*F*_6_,_86_ = 0.18, *P* = 0.98). The 32 °C shift was significant in all four naive age groups (1.2–3.9 days) and in challenged mosquitoes at days 5, 10 and 15 (2.2–2.7 days; Table 1, Supplementary Table S3 and Supplementary Fig. S1a). All test statistics are listed in Supplementary Table S2. The 30 °C shift was 44% of the 32 °C shift (s.e. 10%), close to the 60% expected for a straight line across temperature (test for curvature, *P* = 0.10), so transcriptomic age rose roughly in proportion to temperature, by about half a day per degree.

**Table 1:**
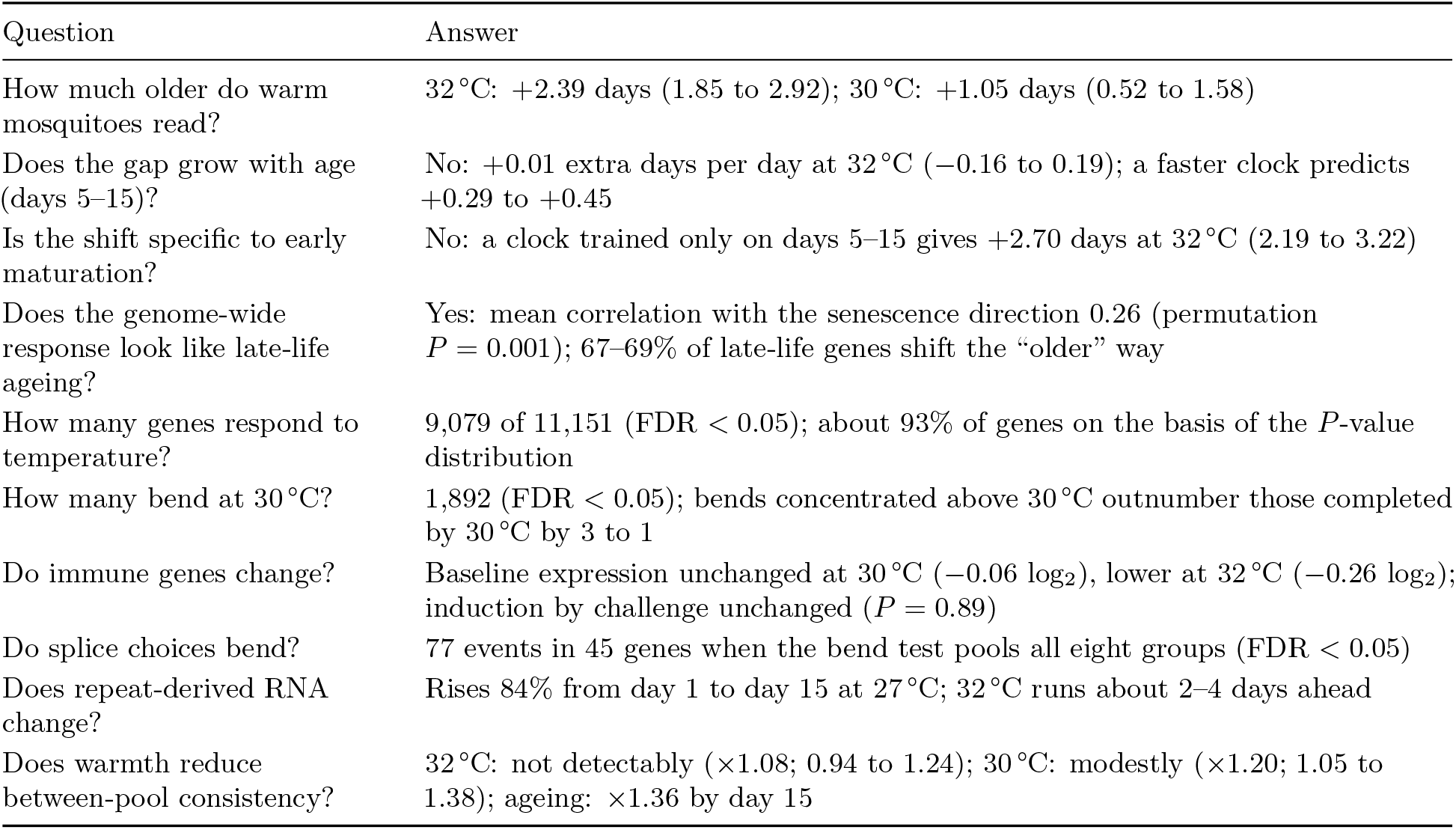
Main results at a glance. Estimates are cohort-adjusted; intervals are 95% confidence intervals. FDR, Benjamini–Hochberg false-discovery rate.

| Question | Answer |
| --- | --- |
| How much older do warm mosquitoes read? | 32 °C: +2.39 days (1.85 to 2.92); 30 °C: +1.05 days (0.52 to 1.58) |
| Does the gap grow with age (days 5–15)? | No: +0.01 extra days per day at 32 °C (−0.16 to 0.19); a faster clock predicts +0.29 to +0.45 |
| Is the shift specific to early maturation? | No: a clock trained only on days 5–15 gives +2.70 days at 32 °C (2.19 to 3.22) |
| Does the genome-wide response look like late-life ageing? | Yes: mean correlation with the senescence direction 0.26 (permutation $P = 0.001$); 67–69% of late-life genes shift the “older” way |
| How many genes respond to temperature? | 9,079 of 11,151 (FDR < 0.05); about 93% of genes on the basis of the $P$-value distribution |
| How many bend at 30 °C? | 1,892 (FDR < 0.05); bends concentrated above 30 °C outnumber those completed by 30 °C by 3 to 1 |
| Do immune genes change? | Baseline expression unchanged at 30 °C (−0.06 log<sub>2</sub>), lower at 32 °C (−0.26 log<sub>2</sub>); induction by challenge unchanged ($P = 0.89$) |
| Do splice choices bend? | 77 events in 45 genes when the bend test pools all eight groups (FDR < 0.05) |
| Does repeat-derived RNA change? | Rises 84% from day 1 to day 15 at 27 °C; 32 °C runs about 2–4 days ahead |
| Does warmth reduce between-pool consistency? | 32 °C: not detectably ($\times 1.08$; 0.94 to 1.24); 30 °C: modestly ($\times 1.20$; 1.05 to 1.38); ageing: $\times 1.36$ by day 15 |

### The gap is set early and then stops growing

If warmth made the clock tick faster, the gap between 32 °C and 27 °C mosquitoes should widen steadily with age. It did not (Fig. 2c). Pooled over immune state, the 32 °C gap was 0.58 days at day 1 (95% CI 0.08–1.08), 2.8 days at day 5, 3.1 days at day 10 and 3.0 days at day 15. Between days 5 and 15, the extra transcriptomic age gained per day of life at 32 °C was 0.01 days (95% CI *−*0.16 to 0.19), and at 30 °C it was 0.10 days (*−*0.07 to 0.28; test for any difference in slope, *F*_2_,_74_ = 0.89, *P* = 0.42). A model in which warmth adds a constant offset fitted the day 5–15 data much better than one in which warmth changes the rate of ageing (Akaike information criterion 43.1 versus 58.0), and adding rate terms to the offset model did not improve it (*F*_4_,_68_ = 0.51, *P* = 0.73).

To judge what a faster clock would look like, we used the thermal-time idea that underlies most insect development models: the rate of a biological process increases in proportion to the temperature above a lower threshold. For any threshold between 10 and 16 °C, a mosquito at 32 °C would gain 0.29–0.45 extra days of transcriptomic age per day relative to one at 27 °C. The observed upper confidence limit, 0.19, excludes this whole range. Put simply, under a faster clock the 32 °C gap of 2.8 days at day 5 would have grown to roughly 6–7 days by day 15; it stayed at 3.0. Warmth therefore shifted the clock forward during the first days of adult life and then left its pace unchanged.

### The shift survives every check we tried

A shift of this kind could, in principle, be an artefact of how the clock was built or how the reads were processed. We therefore rebuilt the clock 28 more times, varying the number of genes (50–2,000), the number of principal components (2–5) and the method (principal-component regression or elastic net on all 11,151 genes), and training on naive and challenged mosquitoes together. Every construction read age accurately (mean absolute error 0.77–1.05 days) and every one gave a clear 32 °C shift, ranging from 1.85 to 3.08 days (Fig. 2d and Supplementary Fig. S1b). The shift was essentially unchanged when gene counts were derived directly from the raw sequencing reads with an independent alignment and counting pipeline (2.33 days), when seven library-quality metrics were added as covariates (2.30 days), and when sequencing batch replaced cohort as the blocking factor (2.28 days). Library-quality metrics differed little between temperatures (Supplementary Fig. S2d). The 30 °C shift was smaller and more sensitive to these choices (0.77–1.95 days) but always positive.

### Warmth advances the late-life programme, not the whole of ageing

Adult mosquitoes change in two broad phases. In the first days after emergence they mature: the cuticle hardens, flight muscles and the gut finish developing and the fat body is restocked. Later, they senesce. In naive and challenged 27 °C mosquitoes, 8,714 genes changed by more than 0.05 log_2_ units per day between days 1 and 5, but only 1,176 did so between days 5 and 15 (Fig. 3a). Because the clock is trained on both phases, an older clock reading could in principle reflect faster maturation alone.

**Figure 3:**
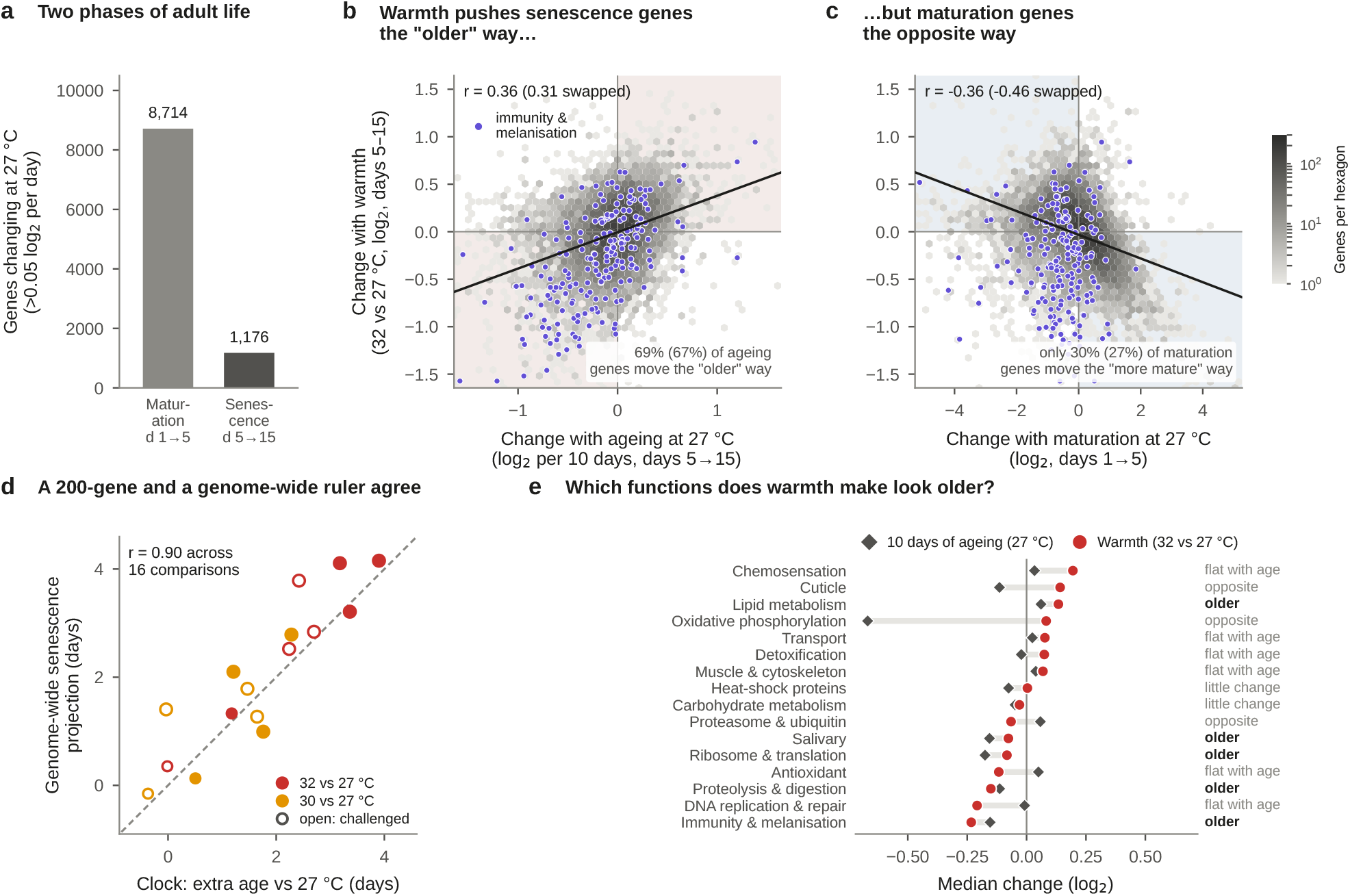
Warmth advances the late-life programme but runs counter to early maturation. **a**, Many more genes change during maturation (days 1*→* 5) than during senescence (days 5*→* 15) at 27 °C. **b**, Each hexagon bins genes by their change during senescence at 27 °C (*x*) and their response to warmth in days 5–15 (*y*). Shaded quadrants are where warmth moves a gene in the same direction as ageing. Ageing directions come from challenged mosquitoes and warmth responses from naive mosquitoes, so no sample contributes to both; values with the roles swapped are given in brackets. Purple points are genes annotated with immune or melanisation functions. **c**, As **b**, for the maturation direction; shaded quadrants are where warmth opposes maturation. **d**, For each of the 16 temperature comparisons, the clock’s estimate of extra age (*x*) against the genome-wide projection of the warmth response onto the senescence direction (*y*), with each group’s own samples excluded from the direction. Filled, naive; open, challenged. **e**, Median change with 10 days of senescence at 27 °C (diamonds) and with warmth (circles) for functional groups with at least 20 genes. Labels mark groups in which both changes exceed 0.05 log_2_ and agree (“older”) or disagree (“opposite”), groups whose change with ageing is below 0.05 log_2_ (“flat with age”), and groups whose change with warmth is below 0.05 log_2_ (“little change”).

It did not. A clock trained only on days 5–15 still read 32 °C mosquitoes as 2.70 days older (95% CI 2.19–3.22) and 30 °C mosquitoes as 1.76 days older (1.25–2.27), again with no change in pace (*−*0.01 extra days per day at 32 °C; *−*0.17 to 0.15). We then asked the question without any clock, using every gene. For each gene we measured its direction of change during senescence (days 5*→*15 at 27 °C) and during maturation (days 1*→*5 at 27 °C), and compared these with its response to warmth in days 5–15. To avoid circularity, the age directions were estimated in one immune state and the warmth response in the other. Warmth pushed late-life genes in the “older” direction: the correlation between the two was 0.36 (0.31 with the roles of the immune states swapped), and 67–69% of genes that change in late life shifted the way ageing would move them (Fig. 3b). In contrast, warmth pushed maturation genes *against* their maturation direction (correlation *−*0.36 and *−*0.46; only 27–30% shifted the “more mature” way; Fig. 3c). Repeating this comparison within each age–immune group, with the group’s own samples left out of the age directions, gave a mean correlation with the senescence direction of 0.26 (permutation *P* = 0.001, 999 permutations of temperature labels within groups). Expressed in days of 27 °C senescence, this genome-wide projection agreed closely with the 200-gene clock across the 16 temperature comparisons (*r* = 0.90; Fig. 3d).

Which functions carried this pattern? Immunity and melanisation genes, digestive proteases and the ribosome and translation machinery all declined during senescence at 27 °C and declined further with warmth (Fig. 3e). Among the largest single contributors to the senescence projection were the prophenoloxidases PPO6 and PPO9, the enzymes that start melanin synthesis, and the digestive serine protease trypsin 2. This transcriptional decline matches the earlier and steeper loss of phenoloxidase activity and melanin deposition measured in these mosquitoes at 30 and 32 °C [3]. Other groups moved the opposite way. Oxidative phosphorylation genes fall steeply during maturation and continue to fall in late life, yet they were higher at 32 °C, as were cuticle, lipid-metabolism, chemosensory and transporter genes. These increases are more easily read as a response to sustained warmth, perhaps reflecting higher metabolic demand, than as ageing. Warmth is thus not simply equivalent to extra time: it advances part of the ageing programme and leaves, or reverses, the rest.

### The middle temperature is rarely a halfway point

Of the 11,151 genes, 9,079 changed with temperature (FDR *<* 0.05), and the distribution of *P* values implied that about 93% of genes respond to some degree. For 1,892 genes, the response was significantly curved: the 30 °C level departed from the straight line joining 27 and 32 °C somewhere in the design (FDR *<* 0.05; Fig. 4a). An independent analysis with limma-voom [7, 8] recovered 97% of the temperature-responsive genes and 80% of the bending genes, and ranked genes almost identically (Spearman *ρ* = 0.98; Supplementary Fig. S2a,b). The *P* -value distribution for the bend test suggested that around 45% of genes bend to some extent, so the 1,892 genes are a conservative subset. The bends were reproducible: 85% of bending genes bent in the same direction in at least six of the eight age–immune groups (Supplementary

**Figure 4:**
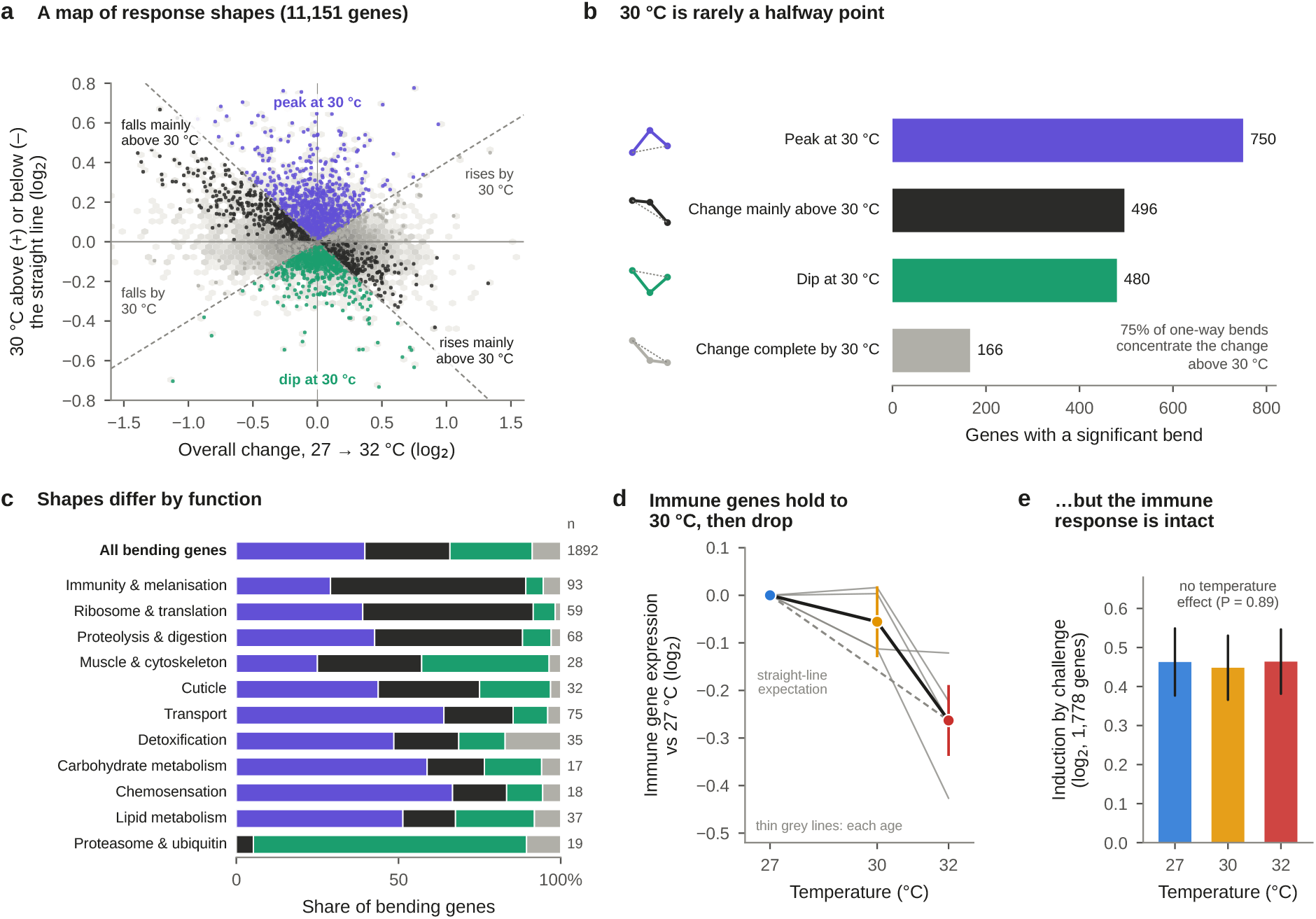
Many responses bend at 30 °C, and immune genes change mainly above it. **a**, Each gene’s overall change from 27 to 32 °C (*x*) against how far its 30 °C level sits above or below the straight line joining 27 and 32 °C (*y*), using cohort-adjusted means pooled over age and immune state. Grey hexagons, all genes; coloured points, the 1,892 genes with a significant bend. Dashed lines divide the four shapes in **b. b**, Bending genes by shape. “Change mainly above 30 °C” means that per degree, the step from 30 to 32 °C is larger than the step from 27 to 30 °C; “complete by 30 °C” means the reverse. **c**, Shapes by functional group (groups with at least 15 bending genes). **d**, Average expression of 204 immune and melanisation genes in naive mosquitoes relative to 27 °C (thick line, cohort- and age-adjusted mean *±* 95% CI; thin lines, each age). **e**, Strength of the response to immune challenge (average change in 1,778 challenge-induced genes) at each temperature (*±* 95% CI).

Fig. S2c).

We classified bending genes into four shapes (Fig. 4b). A peak at 30 °C (750 genes) or a dip at 30 °C (480 genes) means the middle temperature lies outside the range of the other two. The remaining genes changed in one direction, but unevenly. In 496 of them the change was concentrated above 30 °C, and in only 166 was it largely complete by 30 °C (*P* = 5 *×* 10^*-39*^), so when a one-way response bent, the last two degrees usually mattered most. Shapes differed between functions (Fig. 4c). Immune, ribosomal and digestive genes were dominated by changes concentrated above 30 °C, proteasome genes and many muscle genes dipped at 30 °C, and transporter and chemosensory genes tended to peak there.

### Immune genes hold to 30 °C, then fall; the response to challenge is intact

The immune genes illustrate why the middle temperature matters. Averaged across 204 genes annotated with immune or melanisation functions, expression in naive mosquitoes at 30 °C was indistinguishable from 27 °C (*−*0.06 log_2_ units; 95% CI *−*0.13 to 0.02), but at 32 °C it was 0.26 log_2_ units lower (*−*0.34 to *−*0.19; Fig. 4d), about a 17% reduction. The 30 °C level sat 0.10 log_2_ units above the straight-line expectation (*P* = 0.002), so the decline happened mainly between 30 and 32 °C. A two-temperature comparison of 27 and 32 °C would have suggested a gradual decline instead. Strikingly, the capacity to respond to infection signals was untouched. Heat-killed bacteria induced 1,778 genes, and the strength of that induction was the same at all three temperatures (0.46, 0.45 and 0.46 log_2_ units; temperature *×* challenge interaction *F*_8_,_86_ = 0.45, *P* = 0.89; Fig. 4e). Warmth lowered the baseline from which the immune response starts, not the size of the response.

### Splice choices and isoform use also bend

Gene abundance is only one layer of regulation; a gene can also change which exons it includes and which of its transcript isoforms it makes. Splicing changed widely with temperature. Across the 24 pairwise comparisons, 50,732 comparison–event instances, representing 21,899 distinct events in 2,569 genes, differed at FDR *<* 0.05 with a change in exon inclusion of at least 10 percentage points. The fraction of tested events that differed scaled with the size of the temperature step: 2.1% for 30 versus 32 °C, 2.2% for 27 versus 30 °C and 3.4% for 27 versus 32 °C (Fig. 5a).

**Figure 5:**
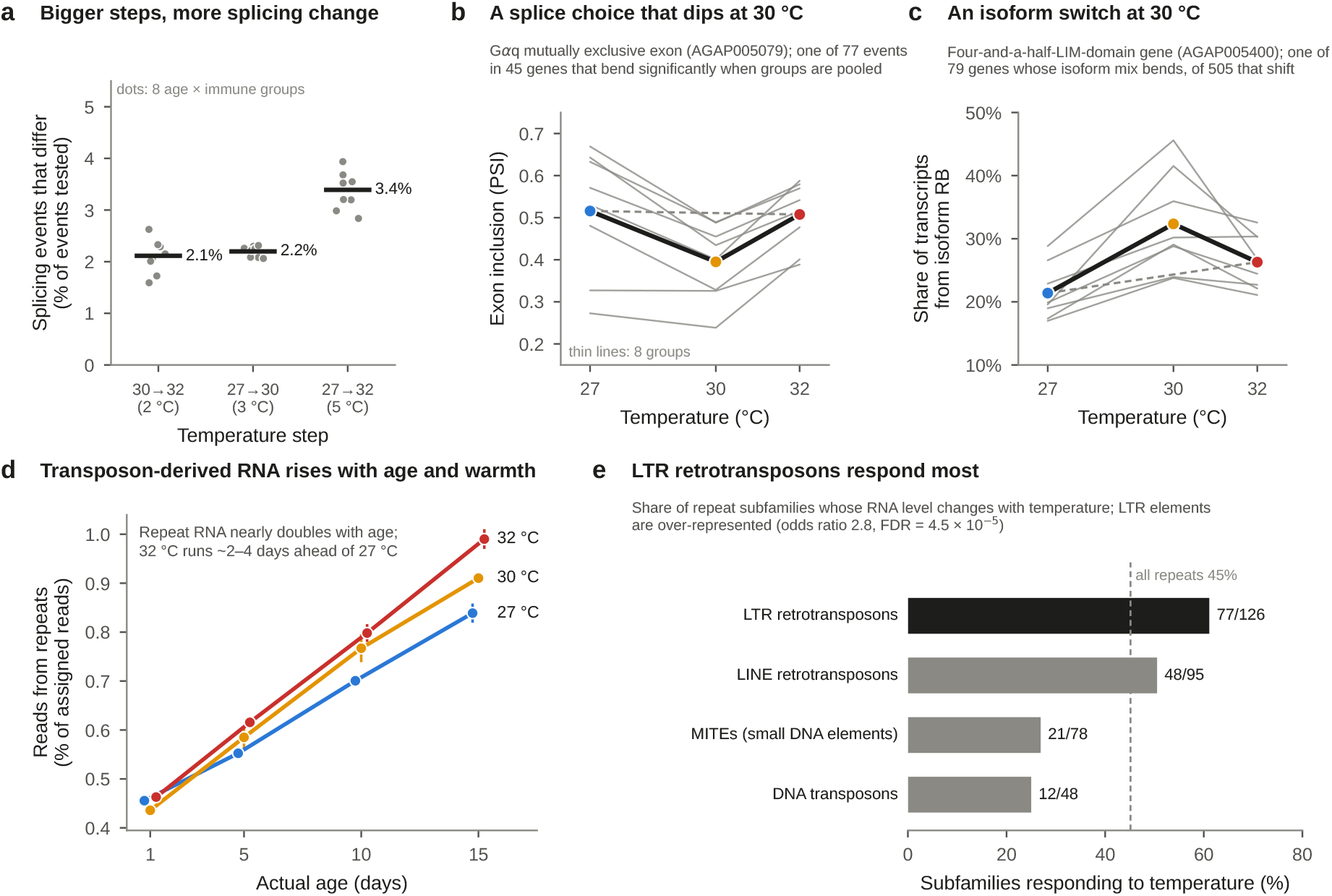
RNA processing and repeat-derived RNA. **a**, Percentage of tested splicing events that differ between temperatures (FDR *<* 0.05, change in exon inclusion 10 *≥*percentage points), by size of the temperature step. Dots, eight age–immune groups; bars, overall. **b**, Exon inclusion (percent spliced in, PSI) of a mutually exclusive exon in the G*α*q gene (AGAP005079), the strongest of 77 bending events. Thin lines, eight age–immune groups; thick line, their mean; dashed line, straight-line expectation. **c**, Share of transcripts from isoform RB of a four-and-a-half-LIM-domain gene (AGAP005400), the strongest of 79 genes with bending isoform use. **d**, Percentage of assigned reads derived from annotated repeats (mostly transposable elements), by age and temperature (mean *±* s.e.; immune states pooled). **e**, Share of repeat subfamilies in each class whose RNA level changed with temperature (FDR *<* 0.05). Dashed line, all repeats.

Detecting bends in splicing requires more statistical power than detecting differences, because exon-inclusion estimates from 4–5 pools are noisy. When the bend test was run separately in each age–immune group, no event survived correction across all 316,518 group–event tests. We therefore pooled all eight groups into a single model with a common bend term, allowing each group its own baseline and slope (Methods). This more powerful test identified 77 events in 45 genes that bent significantly (FDR *<* 0.05, among 14,742 adequately covered events), mostly skipped exons (43) and mutually exclusive exons (13; Supplementary Table S6). In the clearest case, a mutually exclusive exon pair in the G-protein subunit G*α*q gene dipped at 30 °C in seven of eight groups and recovered at 32 °C (Fig. 5b). Seventeen of the 45 genes also bent at the level of overall expression. Isoform use, analysed separately with transcript-level quantification [9, 10], shifted with temperature in 505 of 1,252 multi-isoform genes and bent in 79. Twenty-four genes bent in overall expression, had at least one significant pairwise splicing event and bent in isoform use (Supplementary Table S7); because these layers are not independent, we treat this list as a ranking for follow-up rather than as evidence of shared mechanisms. In one example, a four-and-a-half-LIM-domain gene shifted its output towards isoform RB at 30 °C in seven of eight groups (Fig. 5c).

### Repeat-derived RNA rises with age and runs ahead at 32 °C

About 0.4–1% of reads came from repeats, mostly transposable elements. This fraction is a recognised marker of ageing in insects, in which the silencing of transposable elements loosens with age [11, 12]. In these mosquitoes it nearly doubled with age, from 0.46% at day 1 to 0.84% at day 15 at 27 °C (Fig. 5d). At day 1 the three temperatures were indistinguishable, but by day 15 the fraction was 0.91% at 30 °C and 0.99% at 32 °C. Measured against the age trend, 32 °C mosquitoes ran the equivalent of 2.1–4.2 days ahead of 27 °C mosquitoes, depending on whether the fraction was modelled on the logit scale or as repeat reads relative to gene-based library size, and whether library-quality covariates were included. That is the same order as the clock shift. Across all ages warmth also steepened the age trend (slope ratio 1.27 at 32 °C, *P* = 0.003), but this was driven by divergence after day 1. Within days 5–15 alone, the slopes did not differ detectably (ratio 1.19, *P* = 0.34), so the repeat data cannot separate a shift from a change of pace.

Of 350 repeat subfamilies with enough reads, 158 changed with temperature (FDR *<* 0.05) and 7 bent (Supplementary Fig. S3 and Supplementary Table S5). Long-terminal-repeat retrotransposons responded most often (77 of 126; odds ratio 2.8, FDR = 4.5 *×* 10^*−*5^; Fig. 5e). Within groups, mosquitoes with more repeat RNA also had older clock readings. However, about 60% of this association disappeared once mapping and library-quality metrics were accounted for (variance explained fell from 0.30 to 0.12; Supplementary Fig. S2e), so much of it reflects how reads were assigned rather than biology. These are RNA measurements: they do not show that transposable elements moved or inserted.

### Ageing loosens coordination between pools; 32 °C does not detectably

Ageing is often accompanied by a loss of regulatory precision, so we asked whether warm mosquitoes were also more variable. For each pool, we measured how far its expression profile diverged from the average of the other pools in its group (one minus the rank correlation over the 3,000 most variable genes). Ageing provided a positive control: divergence increased by a factor of 1.42 (95% CI 1.21–1.66) at day 10 and 1.36 (1.16–1.60) at day 15 relative to day 1 (Fig. 6a), showing that the measure can detect a loss of coordination. Against this benchmark, divergence at 32 °C was not detectably higher than at 27 °C (factor 1.08; 0.94– 1.24), whereas 30 °C showed a modest increase (1.20; 1.05–1.38). No individual gene showed a significant change in variability with temperature (0 of 11,151 at FDR *<* 0.05). The older reading at 32 °C is therefore not a by-product of noisier transcription. Because the age trend implies only a 5% increase for 2.4 extra days of age, which lies within the 32 °C confidence interval, this analysis cannot by itself say whether warm mosquitoes are older in this respect too.

**Figure 6:**
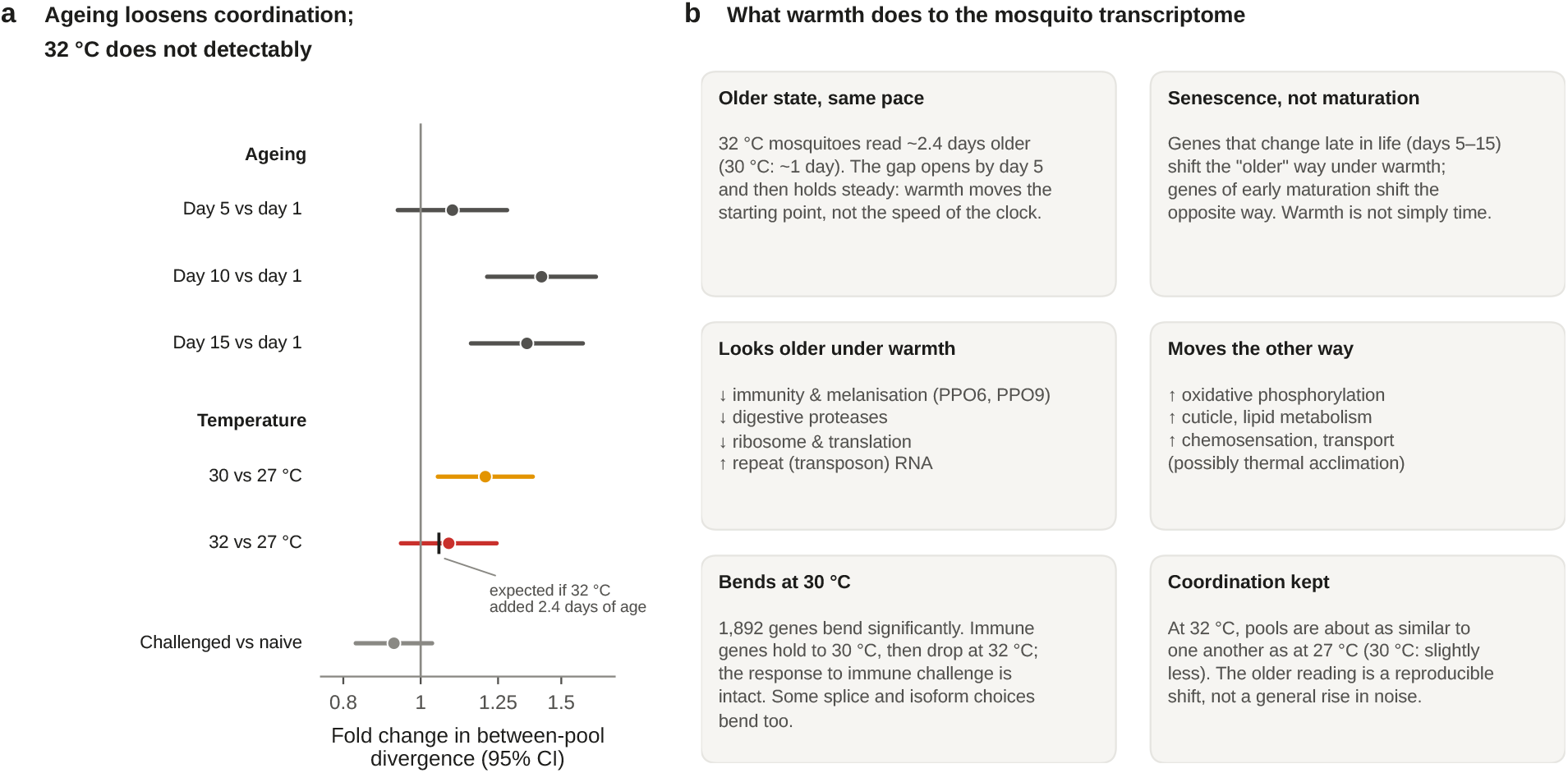
Coordination and synthesis. **a**, Fold change in between-pool divergence (one minus the Spearman correlation between a pool and the average of the other pools in its group, 3,000 most variable genes) for ageing, temperature and immune challenge, from a single cohort-adjusted model on the log scale (95% CI). The black tick on the 32 °C row marks the increase expected if 32 °C mosquitoes were 2.4 days older. **b**, Summary of what warmth does to the mosquito transcriptome in this experiment.

## Discussion

Warmth made mosquitoes look older, but not because their transcriptomic clock ran faster. At 32 °C the clock read mosquitoes as about 2.4 days older than 27 °C mosquitoes of the same age, with a smaller shift at 30 °C. This gap opened during the first five days of adult life and then held steady, which rules out the steadily widening gap expected from a faster clock (Fig. 6b). The finding refines rather than contradicts the conclusion that warmth accelerates senescence in this species [3, 4]. Warm mosquitoes were indeed transcriptionally older, but the difference arose as a step and was not a compounding drift.

Why would warmth produce a step? Because these mosquitoes developed and lived at the same temperature, the step could reflect events during larval and pupal development, during the first days after emergence, or both. The day-1 gap was small (0.6 days), and most of the shift appeared between days 1 and 5, which points to the early adult period. One possibility is that warm mosquitoes settle into a different physiological steady state within a few days: one with higher oxidative-phosphorylation, lipid and transporter expression and lower immune, digestive and translational expression. That state partly overlaps with what ageing does at 27 °C. Another is that early adult life at high temperature causes a burst of wear that is not repeated later. A temperature-switch experiment, moving mosquitoes between 27 and 32 °C at different ages and asking how quickly their clock readings follow, would separate these possibilities in the same way that diet-switch experiments separated the immediate and cumulative effects of dietary restriction [5].

The distinction matters because the two models make different long-term predictions. If warmth increased the pace of ageing, the transcriptional cost of a hot environment would compound over a mosquito’s life and would be largest in exactly the old mosquitoes that transmit malaria. Under a forward shift, a 32 °C mosquito at day 15 is transcriptionally about as old as a 27 °C mosquito at day 18 and stays about three days ahead thereafter. Our data cover only the first 15 days of adult life, and transcriptomic age is not survival, so these results do not show that warming leaves lifespan unchanged. Survival and immune function both decline faster at high temperature [2, 3]. The results do show that, over the ages most relevant to completing the parasite’s development, warming-associated transcriptomic ageing behaved as an offset and not as a change in pace. Models that convert temperature into physiological age by thermal time alone may therefore overstate the transcriptional effect.

Warmth also did not resemble the passage of time in general. It moved genes that change late in life in the ageing direction but moved maturation genes the other way, and it raised oxidative-phosphorylation, cuticle, lipid and chemosensory genes that ageing lowers or leaves alone. The functions that did “age” under warmth are informative. Prophenoloxidases and other melanisation genes, digestive proteases and the translation machinery declined. These declines match the reduced phenoloxidase activity and melanin deposition measured directly at 30 and 32 °C [3], and are consistent with the immune and metabolic themes identified by the source study [4]. Transcriptomic clocks trained on one temperature are therefore best thought of as summarising a particular mixture of processes. Under warming, some of those processes advance and others do not, so a single number such as “2.4 days older” compresses biology that is better described process by process.

The middle temperature changed several conclusions. A comparison of only 27 and 32 °C would suggest that immune gene expression declines steadily with warming, whereas the full series shows that it was maintained at 30 °C and fell between 30 and 32 °C. Across all bending genes, changes concentrated above 30 °C outnumbered changes completed by 30 °C by three to one. This is consistent with the common observation that ectotherm performance curves fall away steeply above an optimum [1, 6], and suggests that the last two degrees in this range carry a disproportionate share of the transcriptional response. Equally important, the induction of challenge-responsive genes was identical at all temperatures, so warmth lowered the immune baseline without blunting the response to a bacterial signal. Whether this lower baseline translates into weaker defence against live pathogens cannot be judged from heat-killed bacteria, but it fits the reduced melanisation seen at high temperature [3].

Two layers beyond gene abundance behaved in line with these conclusions. Splice choices and isoform use changed with temperature and, for a minority of events, bent at 30 °C. The bending events became visible only when the eight age–immune groups were analysed together; testing each group separately lacked the power to find them. Repeat-derived RNA, a marker of ageing in insects [11, 12], rose steeply with age and ran ahead at 32 °C, with long-terminal-repeat retrotransposons responding most often. Its association with clock readings within groups was, however, largely explained by differences in read mapping, a reminder that composition-based measures from the same libraries share technical as well as biological variation.

Finally, the older state at 32 °C was not a disordered one. Between-pool divergence rose with age, confirming that the measure can detect a loss of coordination, but it did not rise detectably at 32 °C. The warm transcriptome was as reproducible as the cool one. Warmth produced a coherent, repeatable change of state rather than a general increase in noise. The analysis was not powerful enough to detect the small increase in divergence that 2.4 extra days of age would imply, so this result constrains how warmth acts rather than whether warm mosquitoes are older.

Several limitations apply. This is a secondary analysis of one experiment on one laboratory strain, and the clock was trained and tested within it (*n* = 18 reference pools). Whole-body pools average across tissues and individuals, so tissue-specific effects may be diluted or hidden. Development and adult life took place at the same temperature, so their contributions cannot be separated. Only three constant temperatures were studied; field temperatures fluctuate, and three points can show that a response bends but not its exact shape. Sampling ended at day 15, before the late-life decline in survival, so whether the offset persists or eventually becomes a change in pace remains open. Functional groups were defined by keywords in gene descriptions, which cover 58% of genes, rather than by curated ontologies, which could not be retrieved in our computing environment. The immune challenge used heat-killed bacteria rather than live infection. Finally, the analyses of pace, ageing phases, response shapes, immune scores and pooled splicing bends were exploratory; we therefore report them with extensive robustness checks and regard them as findings to be confirmed in an independent experiment.

In summary, warmth shifts the malaria mosquito’s transcriptomic clock forward rather than making it run faster. It advances the late-life programme, including melanisation and digestion, while pushing early-maturation and metabolic genes the other way. Many responses, including those of immune genes, change mainly between 30 and 32 °C. Distinguishing shifts from changes of pace, and separating which biological processes are advanced, should help to predict how warming will change the age structure and immune competence of vector populations.

## Methods

### Data and study design

We reanalysed public RNA-sequencing data (NCBI GEO GSE284722; BioProject PRJNA1200107) generated by Martin *et al*. [4, 13]. Female *A. gambiae* (G3 strain) were reared and maintained at 27, 30 or 32 °C and sampled 1, 5, 10 or 15 days after emergence, either naive or 6 h after intrathoracic injection of heat-killed *E. coli*. Each library was a pool of 12–15 whole mosquitoes; stranded poly(A)-selected libraries were sequenced as 150-base paired-end reads. Of 120 libraries, 115 were retained in the source study’s quality-controlled metadata. Four libraries with suspected immune-treatment label swaps and one outlier were excluded before any analysis reported here. Each retained group contained four or five pools. Pools were collected in six experimental cohorts (labelled as replicates in GEO), each spanning several temperatures and ages; cohort was included as a blocking factor in all models. The pool, not the individual mosquito, was the unit of analysis (Supplementary Table S1).

### Gene counts, filtering and transformation

HTSeq gene counts deposited with GSE284722 were matched to samples by identifier; summary rows were removed and gene order was checked across files. Genes with at least 10 reads in at least five pools were retained (11,151 of 13,845). Size factors were estimated by the median-of-ratios method, and counts were transformed with the blind variance-stabilising transformation of DESeq2 [14], using its parametric dispersion trend. For analyses run in Python, the same transformation was implemented following the DESeq2 algorithm; clock predictions obtained with it agreed with those obtained with DESeq2 itself to within 0.01 days on average (correlation *>* 0.9999). Gene names and descriptions were taken from Ensembl Metazoa release 59 (AgamP4) and used only after testing.

### Temperature response and bend tests

For each gene, a negative-binomial generalised linear model (DESeq2) included cohort, age, immune state and their interaction, together with linear (*x* = (*T −* 30)/2) and quadratic (*q* = *x*^2^) temperature terms, each interacting with age and immune state. The bend (nonlinearity) test compared this model with one lacking all quadratic terms (likelihood-ratio test, 8 degrees of freedom); the temperature test compared it with a model lacking all temperature terms (16 degrees of freedom). *P* values were adjusted by the Benjamini–Hochberg procedure [15]. As an independent check, the same nested comparisons were repeated with limma-voom [7, 8], using moderated *F* tests. The proportion of genes with no effect, *π*_0_, was estimated from the *P* -value distribution as the density above *λ* = 0.5 [16].

To describe shapes, we computed cohort-adjusted mean expression on the variance-stabilised scale for each temperature, pooled over age and immune state. Let *d*_1_ be the change from 27 to 30 °C and *d*_2_ the change from 30 to 32 °C. The departure of 30 °C from the straight line joining 27 and 32 °C is

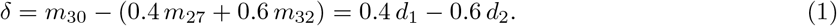

A gene was classed as peaking at 30 °C if *d*_1_ *>* 0 and *d*_2_ *<* 0, dipping if *d*_1_ *<* 0 and *d*_2_ *>* 0, and otherwise as changing mainly above 30 °C if *δ* and the overall change *d*_1_ + *d*_2_ had opposite signs (that is, |*d*_2_|/2 *>* |*d*_1_|/3) or completing its change by 30 °C if they had the same sign. Because *δ* is a contrast of independent means, measurement noise does not favour either sign. Consistency across groups was assessed from the sign of *δ* computed separately in each of the eight age–immune groups.

### Transcriptomic clock

Within each of six folds, one cohort was held out. Using naive 27 °C pools from the remaining cohorts, genes with standard deviation above 0.1 were ranked by the absolute Spearman correlation between expression and age, and the top 200 were standardised and reduced to three principal components. Age in days was regressed on the component scores. All of these steps used training data only, and the fitted transformation was applied to every pool in the held-out cohort, giving one out-of-fold prediction per pool. Accuracy was assessed in held-out naive 27 °C pools. Extra transcriptomic age was defined as predicted minus actual age. Temperature effects on extra transcriptomic age were estimated by ordinary least squares with cohort, age, immune state, their interaction and temperature (as a factor). Omnibus and three-way-interaction tests compared nested full-factorial models by *F* tests. Planned 32 °C-versus-27 °C contrasts within each of the eight age–immune groups were fitted with cohort and temperature, and their *P* values were adjusted together. Curvature in the pooled shift was tested by adding a quadratic temperature term to a model with a linear term.

Robustness was assessed with 28 alternative clocks: principal-component regression with 50, 100, 200, 500, 1,000 or 2,000 genes and 2, 3 or 5 components; elastic-net regression [17, 18] on 200, 1,000 or all 11,151 genes with two penalty strengths; a clock trained on naive and challenged 27 °C pools together; and four late-life clocks trained and evaluated on days 5–15 only. The main clock was also applied to gene counts derived directly from the raw reads (see below), with seven library-quality covariates (uniquely mapped, multi-mapped and too-short-unmapped percentages, mismatch rate, duplication rate, GC content and read depth), and with sequencing batch in place of cohort.

### Shift versus pace

For days 5–15, extra transcriptomic age was modelled with cohort, immune state, temperature, age in days and an age *×* immune state term. Pace was tested by adding temperature-specific age slopes (age *×* 30 °C and age *×* 32 °C); the slope coefficients estimate the extra days of transcriptomic age gained per day of life relative to 27 °C. Offset-only, rate-only and combined models were also compared by the Akaike information criterion. Under a thermal-time model, in which a process runs at a rate proportional to *T −T*_0_, a mosquito at temperature *T* would gain (*T − T*_0_)/(27 *− T*_0_) *−* 1 extra days per day relative to 27 °C; this was evaluated for *T*_0_ between 10 and 16 °C. Because clock predictions are slightly compressed towards the mean (slope of predicted on actual age 0.90 in reference pools), we also multiplied this expectation by 0.90 (giving 0.26–0.41 extra days per day at 32 °C); the conclusion was unchanged.

### Maturation and senescence directions

For each gene, a maturation slope (log_2_ change per day between days 1 and 5) and a senescence slope (days 5, 10 and 15) were estimated by least squares in 27 °C pools, adjusting for cohort. The warmth response was the difference between cohort-adjusted 32 °C and 27 °C means, averaged over days 5, 10 and 15. For the gene-wise comparisons in Fig. 3b,c, slopes were estimated in one immune state and warmth responses in the other, so that no pool contributed to both, and the analysis was repeated with the roles swapped. For the per-group analysis (Fig. 3d), slopes were estimated from all 27 °C pools except those of the group being evaluated. The warmth response ∆ in that group was projected onto the slope vector *s* as ∆ *· s*/(*s · s*), which expresses the response in days of 27 °C ageing, and the correlation between ∆ and *s* was recorded. Significance of the mean correlation across the eight 32 °C-versus-27 °C comparisons was assessed by permuting temperature labels within groups (999 permutations).

### Functional groups and immune scores

Seventeen functional groups were defined by regular expressions applied to Ensembl gene symbols and descriptions (Supplementary Table S4); 58% of genes carried a description. Group-level summaries used medians of gene-level estimates. The immune score of a pool was its mean variance-stabilised expression across the 204 genes in the immunity and melanisation group. Effects of temperature on the naive immune score were estimated with cohort and age as covariates, and the departure of 30 °C from the straight line was tested as a linear contrast. Challenge-induced genes were those with a challenge main-effect *t* statistic above 5 in a model with cohort, temperature, age and immune state (1,778 genes). Their mean expression was modelled with the full factorial design, and the temperature *×* challenge terms (including three-way terms; 8 degrees of freedom) were tested by a nested *F* test. Because selection used the challenge main effect, it did not favour any temperature interaction.

### Read processing, splicing and isoform use

All 240 FASTQ files were downloaded from ENA and checked against their MD5 checksums. Reads were trimmed with fastp [19] (automatic adapter detection, Phred 20, minimum length 35) and aligned to AgamP4 with STAR 2.7.11a [20] in two-pass mode. Transcripts were quantified with Salmon [21] against a decoy-aware index with sequence- and GC-bias correction. Pairwise differential splicing was tested with rMATS 4.4.0 [22] on uniquely mapped reads for 30 versus 27, 32 versus 27 and 32 versus 30 °C within each age–immune group (24 comparisons). An event was called when FDR *<* 0.05 and |∆PSI| *≥* 0.10.

To test for bends in splicing, exon inclusion (PSI) was computed per pool from rMATS junction counts, normalised by the inclusion and skipping form lengths, for events present in both the 30-versus-27 and 32-versus-30 °C comparisons. Values with fewer than 10 supporting reads were discarded. Events were analysed when at least four groups had two or more pools at every temperature and PSI varied by at least 0.02 (standard deviation); 14,742 events in 2,971 genes qualified. For each event, a least-squares model with group-specific intercepts and linear temperature slopes plus cohort was compared with the same model plus a single quadratic term shared across groups (*F* test, 1 degree of freedom), and *P* values were adjusted across events. For comparison, the test was also run separately within each age–immune group (316,518 tests).

Isoform use was analysed with tximport [23] and DRIMSeq [9]. Transcripts were retained with at least 5 counts and 1% of their gene’s total in at least five pools, and genes with at least 10 counts in at least five pools. Dirichlet–multinomial models with the full linear-plus-quadratic design were compared with linear-only and no-temperature models by likelihood-ratio tests, and transcript-level confirmation used stageR [10] at 5% overall FDR.

### Repeat-derived RNA

A repeat annotation for AgamP4 was reconstructed from the RepeatMasker (TEfam library) tables of the Ensembl Metazoa release-59 core database (108,243 insertions; 387 subfamilies; six classes). Reads were assigned to genes and repeats jointly with TEcount (TEtranscripts 2.2.4) [24] in multi-mapping mode with expectation–maximisation. The repeat fraction of a pool was the sum of repeat counts divided by the sum of gene and repeat counts. It was analysed on the logit scale and, alternatively, as repeat counts relative to gene-based size factors, with cohort, immune state, temperature and age in days, plus temperature-specific age slopes where stated. Age-equivalent differences were obtained by dividing the temperature coefficient by the age coefficient. Subfamilies with at least 10 reads in at least five pools (350) were tested with the DESeq2 models used for genes, and class enrichment was tested by Fisher’s exact test. The within-group association between repeat fraction and clock reading was estimated after adjusting for the full factorial design, with and without six mapping and library-quality covariates (the seven listed above except read depth).

### Between-pool divergence and gene-wise variability

For each pool, divergence was one minus the Spearman correlation between its profile and the mean profile of the other pools in its group, over the 3,000 genes with the highest variance across all pools (chosen without reference to the design). Log divergence was modelled with cohort, temperature, age and immune state (main effects). Full-factorial omnibus tests were also run. Gene-wise differences in variability were tested with a Brown–Forsythe-type model of absolute deviations from group medians [25], comparing full-factorial and temperature-free models.

### Statistics and reproducibility

All tests were two-sided. FDR control used the Benjamini–Hochberg procedure within the stated families. Confidence intervals are 95% *t*-based intervals unless stated otherwise. No pools were excluded beyond the five removed by the source study, and no technical replicates were treated as independent. Gene-level likelihood-ratio tests, isoform-use tests, rMATS and repeat-subfamily tests were run in R (4.4–4.6) with DESeq2 1.46–1.52, DRIMSeq 1.34.0, stageR 1.28.0 and tximport 1.34.0. The clock, pace, ageing-direction, shape, immune, pooled-splicing, repeat-fraction, divergence and limma-voom analyses were run in Python 3.13 with NumPy 2.5, SciPy 1.18, pandas 3.0 and scikit-learn 1.9. Random operations used fixed seeds. The analyses of pace, ageing direction, response shape, immune scores and pooled splicing bends were exploratory.

## Data availability

Raw reads and count matrices are available from NCBI GEO (GSE284722) and SRA (BioProject PR-JNA1200107). Processed data, statistical result tables and source data for every figure panel are available from Zenodo (doi:10.5281/zenodo.23337400) [26].

## Code availability

All analysis and figure code, comprising the Python analysis scripts and the R and shell pipelines used for read processing, splicing, isoform-use and repeat analyses, is available from the same Zenodo record [26].

## Acknowledgements

We thank the investigators who generated and deposited GSE284722 for making raw reads, count matrices, metadata and source-study code publicly available.

## Author contributions

K.S.: conceptualisation, methodology, formal analysis, data curation, visualisation, writing (original draft, review and editing), project administration. N.G.: investigation, validation, writing (original draft, review and editing). S.B.: investigation, validation, writing (original draft, review and editing).

## Use of generative AI

Generative artificial-intelligence tools were used to assist the authors with code development, figure preparation, and drafting and editing of the text. The authors conceived and directed the study, reviewed and verified all analyses and results, interpreted the findings and approved the final manuscript, and they take full responsibility for its content.

