## Supplementary information for "Warmth shifts rather than speeds transcriptomic ageing in the malaria mosquito *Anopheles gambiae*"

### Contents

**Supplementary Figures** **1**

**Supplementary Tables** **3**

### Supplementary Figures

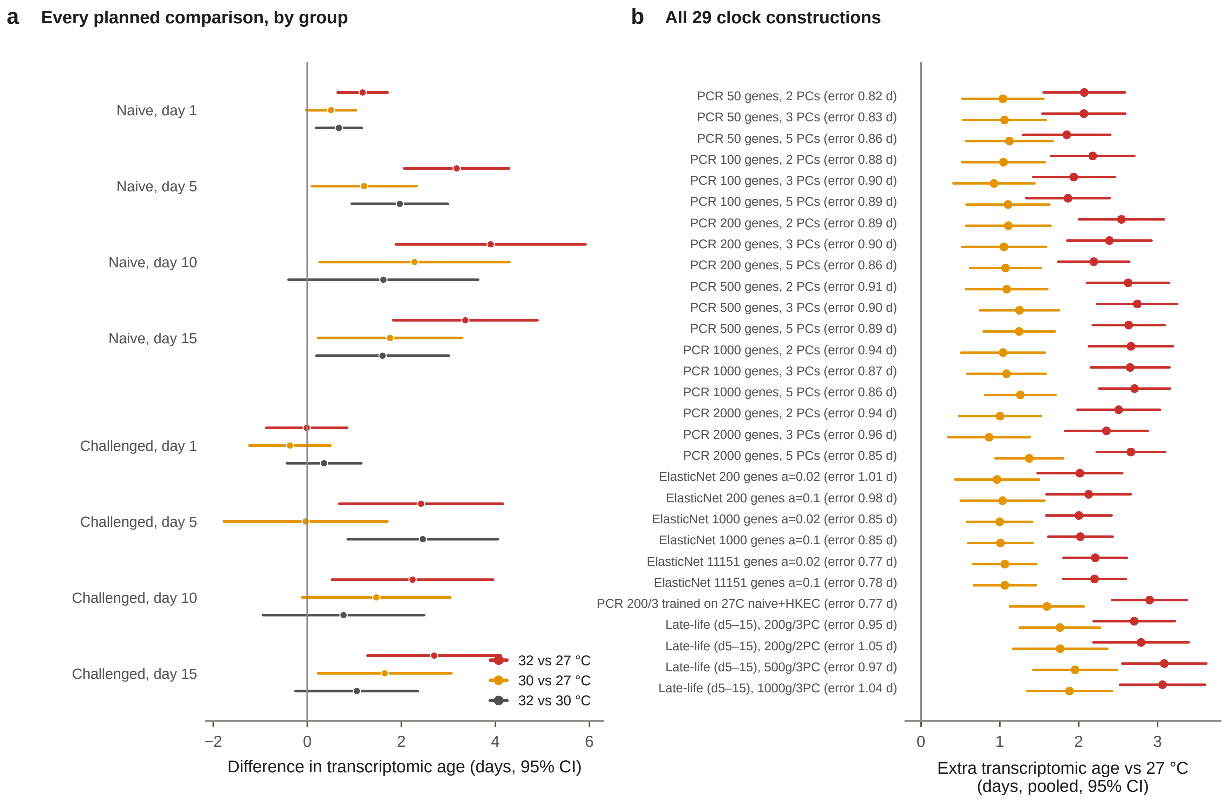


**Figure S1: Clock contrasts by group and under alternative clock constructions.** **a**, Difference in transcriptomic age for each planned temperature comparison within each age–immune group (cohort-adjusted, 95% CI; values in Supplementary Table S3). **b**, Pooled extra transcriptomic age at 32 °C (red) and 30 °C (amber) relative to 27 °C for all 29 clock constructions, with each clock's mean absolute error in held-out naive 27 °C pools (late-life clocks: days 5–15 only). PCR, principal-component regression; PCs, principal components; ElasticNet $a$, penalty strength.

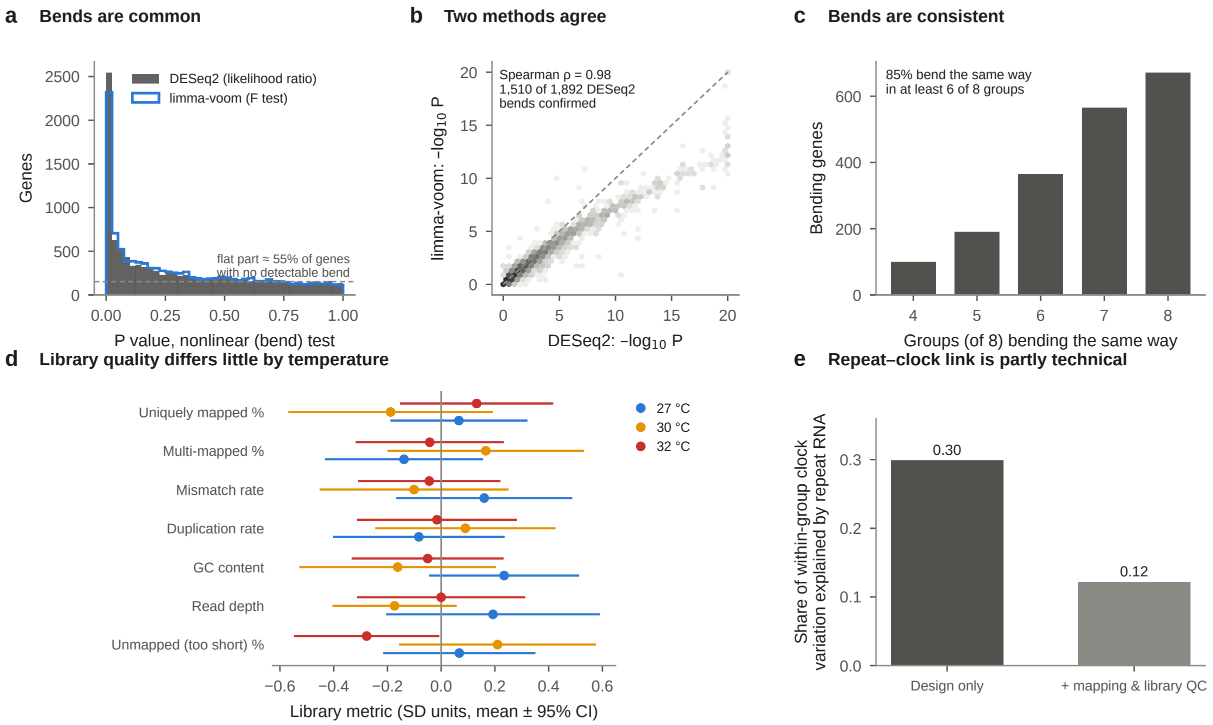


**Figure S2: Statistical and technical cross-checks.** **a**, $P$-value distributions for the bend (nonlinearity) test from DESeq2 and from limma-voom. The dashed line marks the height expected if 55% of genes had no bend ($\pi_0$ estimated at $\lambda = 0.5$). **b**, Agreement between the two methods ($-\log_{10} P$, capped at 20). **c**, Number of bending genes whose 30 °C departure has the same sign in a given number of the eight age–immune groups. **d**, Library-quality metrics by temperature (standardised; mean ± 95% CI). **e**, Share of within-group variation in clock readings explained by the repeat-derived RNA fraction, before and after adjustment for six mapping and library-quality metrics.

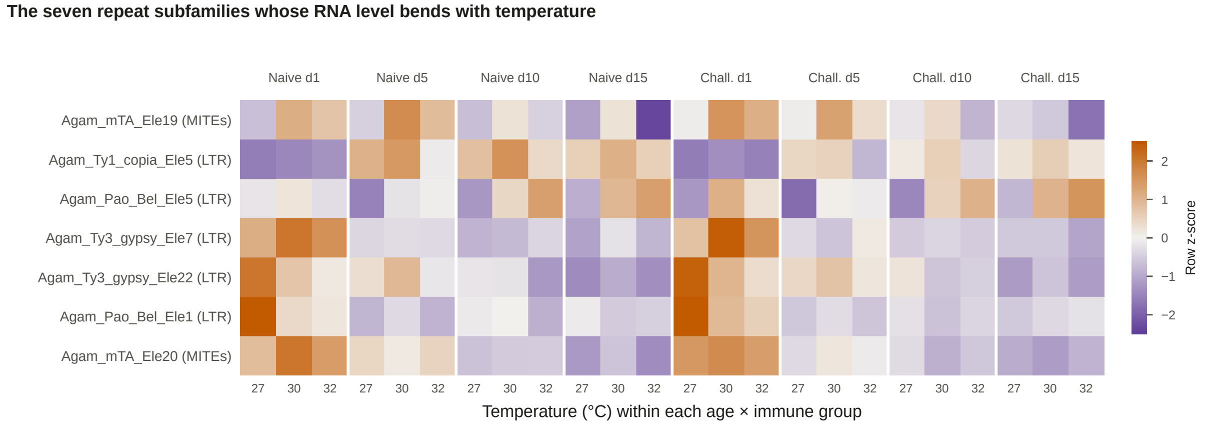


**Figure S3: Repeat subfamilies with a significant bend in RNA level.** Mean normalised expression ($\log_2$) for each temperature within each age–immune group, standardised within rows (purple, below the row mean; orange, above). The scale does not compare absolute abundance between subfamilies. Details in Supplementary Table S5.

### Supplementary Tables

**Table S1: Experimental design.** 115 of 120 libraries were retained by the source study's quality control.

| Factor | Levels |
|---|---|
| Temperature | 27, 30 and 32 °C, constant through development and adult life |
| Adult age | 1, 5, 10 and 15 days after emergence |
| Immune state | Naive; 6 h after injection of heat-killed *Escherichia coli* |
| Pools per group | 4–5 (24 groups); each pool 12–15 whole females |
| Cohorts | Six experimental cohorts (GEO replicate labels), used as a blocking factor |
| Sequencing | Stranded poly(A) RNA-seq, 150-base paired-end reads, Illumina NovaSeq 6000 |
| Source | NCBI GEO GSE284722; BioProject PRJNA1200107 |

**Table S2: Principal statistical results.** Cohort-adjusted throughout. df, degrees of freedom.

| Analysis | Result |
|---|---|
| *Gene expression* | |
| Any temperature effect (DESeq2 LRT, 16 df) | 9,079 of 11,151 genes, FDR $<$ 0.05; limma-voom 8,914 (8,848 shared) |
| Bend at 30 °C (DESeq2 LRT, 8 df) | 1,892 genes, FDR $<$ 0.05; limma-voom 1,527 (1,510 shared); Spearman $\rho = 0.98$ |
| Proportion of genes with no effect ($\pi_0$) | Temperature 0.07; bend 0.55 |
| Bend shapes (bending genes) | Peak 750; dip 480; change mainly above 30 °C 496; complete by 30 °C 166 (binomial $P = 5 \times 10^{-39}$) |
| Consistency of bend direction | 85% of bending genes same sign in $\geq$6 of 8 groups |
| *Transcriptomic clock* | |
| Accuracy, held-out naive 27 °C ($n = 18$) | MAE 0.90 d; RMSE 1.20 d; $R^2 = 0.95$ |
| All temperature terms | $F_{16,86} = 6.89$, $P = 7.9 \times 10^{-10}$ |
| Temperature × age × immune state | $F_{6,86} = 0.18$, $P = 0.98$ |
| Pooled shift, 32 °C vs 27 °C | +2.39 d (1.85 to 2.92), $P = 3.5 \times 10^{-14}$ |
| Pooled shift, 30 °C vs 27 °C | +1.05 d (0.52 to 1.58), $P = 1.6 \times 10^{-4}$ |
| Curvature of the shift across temperature | $F_{1,100} = 2.82$, $P = 0.096$; 30 °C shift 44% (s.e. 10%) of 32 °C shift |
| Pace, days 5–15 (extra days per day) | 32 °C: 0.01 (−0.16 to 0.19); 30 °C: 0.10 (−0.07 to 0.28); $F_{2,74} = 0.89$, $P = 0.42$ |
| Offset versus rate models, days 5–15 | AIC 43.1 vs 58.0; rate beyond offset $F_{4,68} = 0.51$, $P = 0.73$; offset beyond rate $F_{4,68} = 3.77$, $P = 0.008$ |
| Late-life clock (days 5–15) | 32 °C: +2.70 d (2.19 to 3.22); 30 °C: +1.76 d (1.25 to 2.27); pace at 32 °C −0.01 (−0.17 to 0.15) |
| *Ageing directions* | |
| Gene-wise warmth vs senescence direction (disjoint samples) | $r = 0.36$ and 0.31; 69% and 67% of late-life genes shift "older" |
| Gene-wise warmth vs maturation direction (disjoint samples) | $r = -0.36$ and $-0.46$; 30% and 27% shift "more mature" |
| Per-group held-out projection (8 comparisons, 32 °C) | Mean $r$ senescence 0.26 (permutation $P = 0.001$); maturation −0.19 ($P = 0.06$) |
| Clock vs genome-wide senescence projection | $r = 0.90$ across 16 comparisons |
| *Immunity* | |
| Naive immune score (204 genes) | 30 °C: −0.06 (−0.13 to 0.02); 32 °C: −0.26 (−0.34 to −0.19) $\log_2$; 30 °C departure +0.10 (s.e. 0.03), $P = 0.002$ |
| Induction by challenge (1,778 genes) | 0.46, 0.45, 0.46 $\log_2$ at 27, 30, 32 °C; temperature × challenge $F_{8,86} = 0.45$, $P = 0.89$ |
| *RNA processing* | |
| Pairwise splicing (rMATS, 24 comparisons) | 50,732 instances; 21,899 events; 2,569 genes; 1,973 intron-retention instances |
| Splicing bends, per-group tests | 0 of 316,518 tests at global FDR $<$ 0.05 |

| Analysis | Result |
|---|---|
| Splicing bends, pooled test | 77 of 14,742 events (45 genes) at FDR $< 0.05$; $\pi_0 = 0.89$ |
| Isoform use (DRIMSeq) | Temperature: 505 of 1,252 genes; bend: 79 genes (FDR $< 0.05$) |
| *Repeat-derived RNA* | |
| Subfamily tests (DESeq2) | Temperature 158 of 350; bend 7; LTR enrichment OR 2.77, FDR $4.5 \times 10^{-5}$ |
| Repeat fraction, temperature (logit) | Omnibus $F_{16,86} = 3.52$, $P = 7.7 \times 10^{-5}$; quadratic $F_{8,86} = 0.73$, $P = 0.67$ |
| Age-equivalent lead at 32 °C | 2.1–4.2 days across five model specifications |
| Temperature-specific age slopes | All ages: ratio 1.27, $P = 0.003$; days 5–15: ratio 1.19, $P = 0.34$ |
| Within-group repeat–clock association | Partial $R^2$ 0.30 ($P = 4 \times 10^{-8}$); with QC covariates 0.12 ($P = 0.001$) |
| *Between-pool divergence* | |
| Full-factorial temperature terms | $F_{16,86} = 1.18$, $P = 0.30$ |
| Main-effects model (log scale) | Age $F_{3,103} = 8.84$, $P = 3 \times 10^{-5}$; temperature $F_{2,103} = 3.70$, $P = 0.028$ |
| Fold changes | Day 10: 1.42 (1.21–1.66); day 15: 1.36 (1.16–1.60); 30 °C: 1.20 (1.05–1.38); 32 °C: 1.08 (0.94–1.24) |
| Gene-wise variability | 0 of 11,151 genes at FDR $< 0.05$ |

**Table S3: Differences in transcriptomic age within each age–immune group.** Estimate in days (standard error); FDR-adjusted $P$ across the eight groups for each comparison.

| Immune state | Day | 30 vs 27 °C | 32 vs 27 °C | 32 vs 30 °C |
|---|---|---|---|---|
| Naive | 1 | 0.51 (0.22); 0.084 | 1.18 (0.22); 0.003 | 0.67 (0.21); 0.037 |
| Naive | 5 | 1.21 (0.47); 0.074 | 3.18 (0.47); 0.002 | 1.97 (0.43); 0.021 |
| Naive | 10 | 2.28 (0.87); 0.074 | 3.90 (0.87); 0.004 | 1.62 (0.87); 0.135 |
| Naive | 15 | 1.76 (0.65); 0.074 | 3.36 (0.65); 0.003 | 1.60 (0.60); 0.062 |
| Challenged | 1 | -0.37 (0.36); 0.393 | -0.01 (0.36); 0.972 | 0.36 (0.33); 0.323 |
| Challenged | 5 | -0.04 (0.74); 0.963 | 2.42 (0.74); 0.018 | 2.46 (0.67); 0.033 |
| Challenged | 10 | 1.47 (0.67); 0.084 | 2.24 (0.73); 0.020 | 0.77 (0.73); 0.323 |
| Challenged | 15 | 1.65 (0.60); 0.074 | 2.70 (0.60); 0.004 | 1.05 (0.55); 0.135 |

**Table S4: Keyword definitions of functional groups.** Case-insensitive regular expressions applied to Ensembl Metazoa release-59 gene symbols and descriptions (\b, word boundary; \d, digit). 58% of the 11,151 genes carried a description.

<table>
<tr><th>Group</th><th>Pattern</th></tr>
<tr><td>Heat-shock proteins & chaperones</td><td>heat shock | \bhsp | dnaj | chaperon | crystallin | hsc70 | t-complex protein</td></tr>
<tr><td>Immunity & melanisation</td><td>\bclip | defensin | cecropin | gambicin | lysozyme | leucine-rich immune | thioester | peptidoglycan recognition | \btep\d | \bapl1 | serpin | c-type lectin | fibrinogen | phenoloxidase | \bppo | toll | \blrim | galectin | attacin | antimicrobial</td></tr>
<tr><td>Cuticle</td><td>cuticular | cuticle | \bcpr\d | chitin</td></tr>
<tr><td>Detoxification</td><td>cytochrome p450 | \bcyp\d | glutathione s-transferase | \bgst | carboxylesterase | \bcoe | udp-glucuronosyl | \bugt</td></tr>
<tr><td>Oxidative phosphorylation</td><td>nadh dehydrogenase | nadh-ubiquinone | cytochrome c oxidase | atp synthase | ubiquinol | cytochrome b-c1 | succinate dehydrogenase | cytochrome c1</td></tr>
<tr><td>Ribosome & translation</td><td>ribosomal protein | \brp[ls]\d | translation initiation | elongation factor | eukaryotic translation</td></tr>
<tr><td>Proteolysis & digestion</td><td>trypsin | chymotrypsin | carboxypeptidase | aminopeptidase | serine protease | peptidase</td></tr>
<tr><td>Lipid metabolism</td><td>lipase | fatty acid | acyl-coa | lipophorin | elongation of very long | desaturase | lipid</td></tr>
<tr><td>Carbohydrate metabolism</td><td>glucosidase | trehalase | maltase | glycogen | hexokinase | phosphofructokinase | pyruvate kinase | aldolase | enolase | glucose</td></tr>
<tr><td>DNA replication & repair</td><td>dna repair | dna polymerase | \brad\d | minichromosome | dna replication | dna ligase | mismatch repair | excision repair | recq | dna helicase</td></tr>
<tr><td>Reproduction (vitellogenesis)</td><td>vitellogenin | vitelline | chorion | vitellogenic</td></tr>
<tr><td>Chemosensation</td><td>odorant | gustatory | olfactory | chemosensory | ionotropic receptor</td></tr>
<tr><td>Salivary</td><td>salivary | \bd7 | apyrase</td></tr>
<tr><td>Muscle & cytoskeleton</td><td>myosin | troponin | tropomyosin | titin | actin | paramyosin | flightin | muscle</td></tr>
<tr><td>Proteasome & ubiquitin</td><td>proteasome | ubiquitin</td></tr>
<tr><td>Antioxidant</td><td>catalase | superoxide dismutase | peroxiredoxin | thioredoxin | glutathione peroxidase</td></tr>
<tr><td>Transport (solute carriers/ABC)</td><td>solute carrier | \bslc | abc transporter | atp-binding cassette | transporter</td></tr>
</table>

**Table S5: Repeat subfamilies with a significant bend.** The group shown is the one with the largest departure of 30 °C from the straight line; shapes are classified as in the main text (Methods).

| Subfamily | Family | Class | FDR | Strongest group | Shape there |
|---|---|---|---|---|---|
| Agam_mTA_Ele19 | mTA | MITEs | 2.3e-10 | 15 d, naive | peak at 30 °C |
| Agam_Ty1_copia_Ele5 | Ty1_copia | LTR | 9.1e-09 | 5 d, naive | peak at 30 °C |
| Agam_Pao_Bel_Ele5 | Pao_Bel | LTR | 4.5e-03 | 1 d, challenged | peak at 30 °C |
| Agam_Ty3_gypsy_Ele7 | Ty3_gypsy | LTR | 4.5e-03 | 1 d, challenged | peak at 30 °C |
| Agam_Ty3_gypsy_Ele22 | Ty3_gypsy | LTR | 8.6e-03 | 5 d, naive | peak at 30 °C |
| Agam_Pao_Bel_Ele1 | Pao_Bel | LTR | 1.7e-02 | 1 d, naive | falls by 30 °C |
| Agam_mTA_Ele20 | mTA | MITEs | 3.7e-02 | 1 d, naive | peak at 30 °C |

**Table S6: The 25 strongest splicing events with a significant bend in the pooled test** (of 77 at FDR < 0.05). Common quadratic coefficient: positive means 30 °C inclusion lies below the straight line (dip), negative above it (peak), in PSI units per $(2\,°C)^2$. SE, skipped exon; MXE, mutually exclusive exons; A5SS/A3SS, alternative $5'/3'$ splice site; RI, retained intron.

| Type | Gene | Description | Location | Groups | Coefficient | FDR |
|---|---|---|---|---|---|---|
| MXE | AGAP005079 | guanine nucleotide-binding protein G(q) alpha su | chr2L:9742779 | 8 | +0.077 | 1.9e-09 |
| A5SS | AGAP001023 | myofilin variant B | chrX:19660857 | 8 | +0.013 | 6.4e-09 |
| SE | AGAP005079 | guanine nucleotide-binding protein G(q) alpha su | chr2L:9743634 | 8 | -0.067 | 2.9e-08 |
| SE | AGAP006366 | 2-oxoglutarate dehydrogenase E1 component | chr2L:30518178 | 8 | -0.012 | 4.0e-08 |
| SE | AGAP005079 | guanine nucleotide-binding protein G(q) alpha su | chr2L:9742779 | 8 | +0.061 | 9.2e-08 |
| MXE | AGAP005079 | guanine nucleotide-binding protein G(q) alpha su | chr2L:9745001 | 8 | -0.059 | 9.2e-08 |
| MXE | AGAP005079 | guanine nucleotide-binding protein G(q) alpha su | chr2L:9745009 | 5 | -0.086 | 4.9e-07 |
| MXE | AGAP005079 | guanine nucleotide-binding protein G(q) alpha su | chr2L:9745001 | 8 | -0.064 | 1.2e-06 |
| SE | AGAP002413 | Basic leucine zipper and W2 domain-containing pr | chr2R:21024470 | 8 | -0.019 | 2.1e-06 |
| A3SS | AGAP003939 | | chr2R:46562757 | 5 | +0.017 | 2.9e-06 |
| SE | AGAP003997 | casein kinase 1, gamma | chr2R:47719443 | 8 | -0.021 | 6.6e-06 |
| A3SS | AGAP011099 | myosin III | chr3L:16874573 | 8 | -0.089 | 6.6e-06 |
| RI | AGAP000669 | | chrX:11840619 | 6 | +0.012 | 1.9e-05 |
| SE | AGAP011099 | myosin III | chr3L:16874610 | 8 | -0.100 | 4.9e-05 |
| SE | AGAP002413 | Basic leucine zipper and W2 domain-containing pr | chr2R:21024301 | 8 | -0.025 | 5.5e-05 |
| SE | AGAP002413 | Basic leucine zipper and W2 domain-containing pr | chr2R:21024106 | 8 | -0.025 | 5.5e-05 |
| SE | AGAP005079 | guanine nucleotide-binding protein G(q) alpha su | chr2L:9745001 | 6 | -0.067 | 7.3e-05 |
| SE | AGAP002413 | Basic leucine zipper and W2 domain-containing pr | chr2R:21024103 | 8 | -0.024 | 1.0e-04 |
| SE | AGAP003997 | casein kinase 1, gamma | chr2R:47723580 | 8 | -0.028 | 1.1e-04 |
| SE | AGAP002413 | Basic leucine zipper and W2 domain-containing pr | chr2R:21024459 | 7 | -0.027 | 1.3e-04 |
| MXE | AGAP001936 | phosphatidylinositol phospholipase C, beta | chr2R:12656331 | 8 | +0.039 | 1.9e-04 |
| SE | AGAP004362 | glutamate dehydrogenase (NAD(P)+) | chr2R:55059577 | 8 | +0.041 | 3.0e-04 |
| SE | AGAP007396 | epsin | chr2L:46310757 | 8 | -0.019 | 8.4e-04 |
| MXE | AGAP002858 | Sodium/potassium-transporting ATPase subunit alp | chr2R:28379752 | 8 | +0.015 | 9.5e-04 |
| SE | AGAP005563 | facilitated trehalose transporter Tret1 | chr2L:17077530 | 6 | -0.050 | 1.5e-03 |

**Table S7: Exploratory cross-layer candidates.** Genes with a significant bend in expression (FDR < 0.05), at least one significant pairwise splicing event, and a significant bend in isoform use. The layers are not independent and the list is a ranking for follow-up, not a set of validated mechanisms.

| Gene | Description | Expression bend FDR | Min. splicing FDR | Isoform bend FDR |
|---|---|---|---|---|
| AGAP003587 | sodium-independent sulfate anion transporter | 5.4e-19 | 1.5e-08 | 4.4e-03 |
| AGAP002355 | Glutamine synthetase | 8.5e-15 | 1.5e-04 | 1.6e-02 |
| AGAP005563 | facilitated trehalose transporter Tret1 | 1.6e-12 | 1.2e-52 | 3.0e-04 |
| AGAP005795 | sodium-coupled monocarboxylate transporter 1 | 1.5e-09 | 1.9e-06 | 4.6e-02 |
| AGAP009036 | solute carrier family 9 (sodium/hydrogen exchanger), member | 4.7e-08 | 1.8e-12 | 2.1e-02 |
| AGAP000380 | 5'-nucleotidase | 2.6e-05 | 4.2e-07 | 3.6e-03 |
| AGAP029654 | | 3.3e-05 | 2.2e-04 | 2.1e-02 |
| AGAP007753 | | 9.2e-05 | 3.7e-21 | 2.1e-02 |
| AGAP004199 | solute carrier family 5 (sodium-coupled monocarboxylate tran | 3.9e-04 | 2.2e-02 | 4.4e-03 |
| AGAP029666 | | 2.1e-03 | 1.9e-08 | 2.6e-06 |
| AGAP029628 | Myosin-light-chain kinase | 2.2e-03 | 5.5e-54 | 6.1e-03 |
| AGAP001936 | phosphatidylinositol phospholipase C, beta | 2.3e-03 | 2.3e-16 | 4.4e-05 |
| AGAP011676 | uncoupling protein 2, mitochondrial | 3.5e-03 | 5.4e-54 | 9.9e-03 |
| AGAP002274 | | 3.6e-03 | 2.5e-03 | 1.5e-02 |
| AGAP005175 | acetyl-CoA carboxylase / biotin carboxylase | 3.7e-03 | 2.0e-13 | 2.0e-02 |
| AGAP010737 | | 6.0e-03 | 8.5e-04 | 1.1e-03 |
| AGAP002250 | calcium/calmodulin-dependent protein kinase kinase | 8.1e-03 | 7.1e-14 | 4.6e-02 |
| AGAP002324 | mitochondrial sodium/hydrogen exchanger NHA2 | 1.2e-02 | 1.6e-15 | 4.6e-02 |
| AGAP006921 | PID domain-containing protein | 2.4e-02 | 1.0e-68 | 1.3e-02 |
| AGAP007562 | | 3.1e-02 | 2.3e-12 | 9.9e-03 |
| AGAP029610 | | 3.7e-02 | 2.8e-08 | 3.3e-02 |
| AGAP007580 | 60S ribosomal protein L18 | 4.3e-02 | 6.4e-19 | 3.6e-10 |
| AGAP004940 | | 4.5e-02 | 2.3e-11 | 6.4e-05 |
| AGAP003051 | stearoyl-CoA desaturase (delta-9 desaturase) | 4.6e-02 | 6.7e-17 | 4.7e-05 |
